# Stereoscopic RNA Analysis of Transcriptional and Post-transcriptional Regulation

**DOI:** 10.1101/2023.06.22.546182

**Authors:** Seungha Alisa Lee, Hojoong Kwak

## Abstract

Regulation of gene expression takes place at multiple stages of RNA synthesis, processing, and decay. The dynamics of RNA processing is an essential layer of RNA regulation, required for a deeper understanding of gene expression. Here, we present a streamlined analysis combining nascent RNA sequencing and poly(A) tail length sequencing methods, named Stereoscopic Analysis of Transcriptome (STOAT). Using this analysis, we were able to redefine known and unknown transcripts with high precision, and quantitatively assess RNA expression and stability. We also investigated poly(A) tail processing and their linkage to post-transcriptional features in different cell lines and identified the diversity of microRNA networks in RNA stability and opposing effects of HuR-TTP on poly(A) processing among human cell lines. This method can effectively measure promoter activity, RNA synthesis, poly(A) processing, stability, and decay, providing a comprehensive perspective of the dynamic transcriptome as well as discovering diverse transcript isoforms.

**MOTIVATION:** RNA sequencing (RNA-seq) enables transcriptome-wide mapping and quantitative analysis, but it only provides a static view of the transcriptome. Also, due to its lack of standardization between sequencing platforms and read depth, it compromises reproducibility, and the cost remains prohibitive. To improve these limitations, we are presenting the minimal combination of two methods, nascent RNA sequencing and poly(A) tail length sequencing, which can effectively measure promoter activity, RNA synthesis, abundance, poly(A) processing, stability, and decay, and can discover diverse undefined transcript isoforms. This streamlined analysis of transcription rate and polyadenylation status will provide a comprehensive perspective of the dynamic transcriptome.

## INTRODUCTION

Genome-wide transcriptome analysis has become a crucial tool in both understanding gene expression and identifying undefined transcripts^1,2^. Dynamic regulation of gene expression takes place at multiple stages of RNA synthesis, processing, and decay^3^. RNA sequencing (RNA-seq) has been used by providing a comprehensive snapshot of transcript abundance, yet, it has limitations in capturing the dynamics between RNA synthesis and decay^4-6^. RNA-seq can be more powerful when it is combined with rapid perturbations, global inhibition of transcription, or metabolic RNA labeling. However, these approaches require multiple transcriptome measurements in well-defined experimental systems, which can be cost prohibitive^6-8^. Furthermore, while post-transcriptional RNA processing such as deadenylation is coupled to decay, how and whether such processes are linked in a context-specific manner requires scalable tools ^6,7,9-11^.

To address this, we combined nascent RNA (PRO-seq) and poly(A) (TED-seq) measurements^12-15^ to gain a comprehensive view of transcriptional and post-transcriptional processes. PRO-seq (Precision Run On sequencing) measures nascent RNA to identify 5′ transcription start peaks and transcription activity (**Fig 1A**, red and blue). TED-seq (Tail-End Displacement sequencing) measures the abundance of poly(A) RNAs, 3′ cleavage polyadenylation sites (CPS), and poly(A) tail lengths (PAL) simultaneously (**Fig 1A**, orange). We previously developed TED-seq which is cost-efficient and compatible with standard sequencing procedures for higher throughput PAL measurements. Our combination tool of these two methods, called stereoscopic analysis of the transcriptome (STOAT), reveals a deeper view of the dynamics of the transcriptome than ever before.

**Fig 1.**
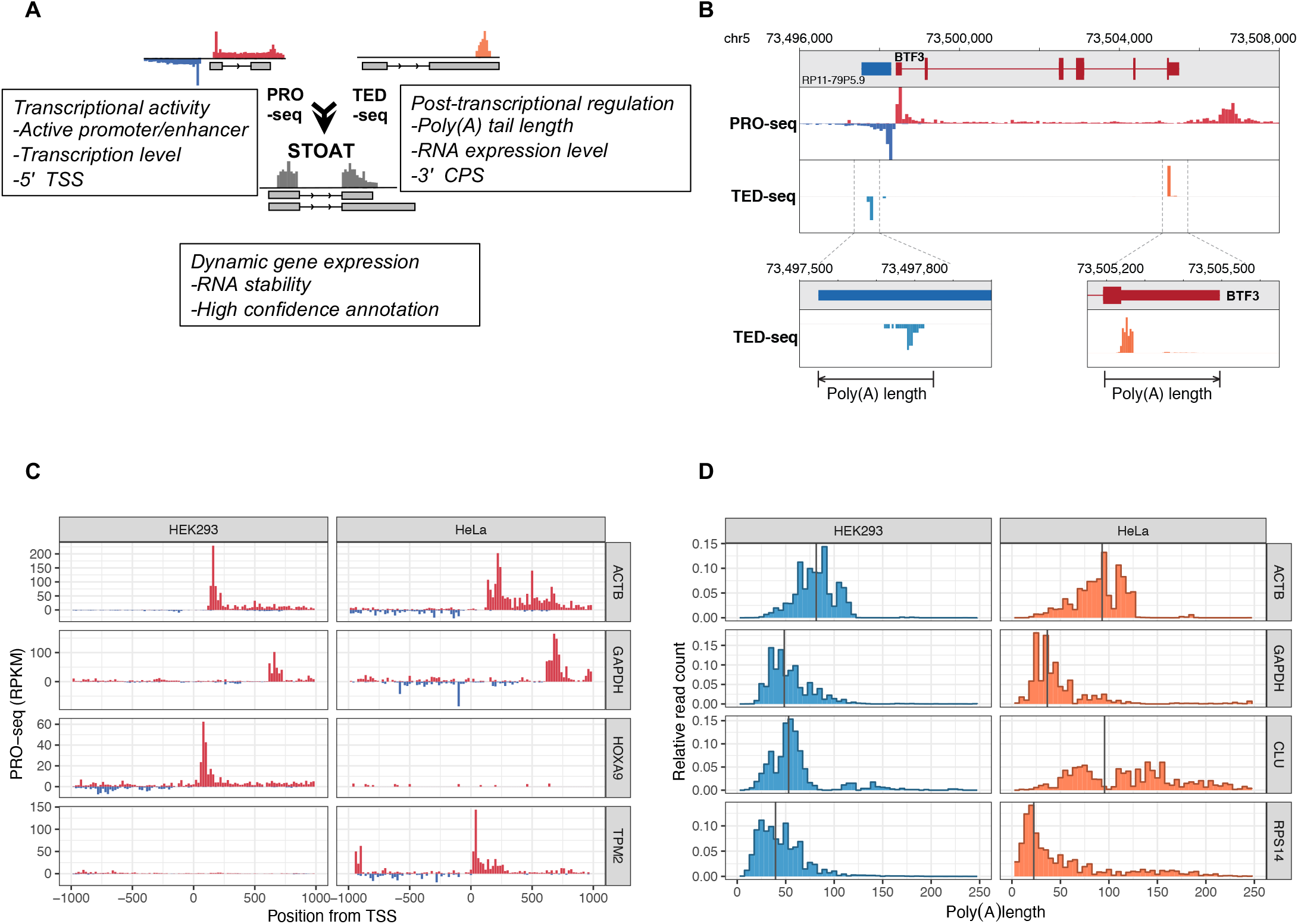
Combined Analysis of Nascent RNA Sequencing and Poly(A) Tail Length Profiling. **A**. Schematic view of STOAT. Combination of Precision Run-On sequencing (PRO-seq) and poly(A) tail length profiling, Tail End-Displacement sequencing (TED-seq), can provide a comprehensive gene regulatory analysis that provides a deeper view of the dynamics of transcriptome. **B**. Example of PRO-seq and TED-seq profiling for *BTF3* gene in integrative genome viewer format. **C**. Examples of PRO-seq expression profiles in different cell type specific regulatory genes, ACTB, GAPDH, HOXA9, and TPM2. **D**. Examples of TED-seq expression profiles in different cell type specific regulatory genes, ACTB, GAPDH, HOXA9, and TPM2.

## RESULTS

### Combined Analysis of Nascent RNA Sequencing and Poly(A) Tail Length Profiling

STOAT is a comprehensive pipeline for adding dimension to RNA dynamics, stability, and regulatory networks, which combines a dual layer of transcriptome data at active transcription and polyadenylation levels (**Fig1A**). Since stable transcripts have higher RNA abundance compared to their transcription level, the ratio between TED-seq and PRO-seq provides an effective measure of steady-state RNA stability. The integration of the two leads to enhance the details of the regulatory steps in gene expression. The characteristic peak patterns of PRO-seq and TED-seq define the 5′ and the 3′ ends of the genes suggesting that this tool can also be used to re-annotate transcription.

In this study, we demonstrated the practical usage of STOAT in a disease-related transcript. For BTF3, a gene associated with non-syndromic deafness in humans, PRO-seq and TED-seq showed a characteristic profile of transcription and polyadenylation in HEK293 cells (**Fig 1B**). Using STOAT, we found that the upstream antisense RNA (uaRNA) of the BTF3 gene was poly-adenylated in addition to its downstream polyadenylated transcripts. TED-seq was able to detect the start of a poly(A) tail containing RNA fragments that were size-selected and showed a read distribution upstream of the 3′ CPS (**Fig 1B**, lower panels). The TED-seq read distribution was equal to the PAL distribution relative to the position 300 bp (library size) upstream of 3′ CPS, and most of the reads in both BTF3 sense transcript and uaRNA showed PALs around 50 bases (**Fig 1B**, lower panels). By using this comprehensive pipeline, we identified global, and cell-type specifically transcribed and polyadenylated transcripts (**Supplementary Text, Fig S1, S2** and **Table S1**). PRO-seq profiles near TSSs showed divergent bidirectional PRO-seq peaks and elongating RNA polymerase densities on the gene body. The housekeeping genes ACTB and GAPDH were transcribed in both HEK293 and HeLa cells, but HOXA9 transcription was HEK293 specific, while TPM2 transcription was HeLa specific (**Fig 1C**). Similarly, TED-seq profiles of ACTB and GAPDH showed comparable PALs. However, PAL of CLU transcript, a gene associated with the coagulation pathway, was longer in HeLa cells, while RPS14 had shorter PAL in HeLa cells (**Fig 1D**). These observations demonstrate that there are cell type-specific regulations of transcription and polyadenylation.

### Reconstitution of RNA Stability and miRNA Network Using STOAT

We also used STOAT to quantitatively reconstitute RNA expression and stability (**Supplementary Text, Fig S3**). When transcripts are in steady state, the ratio between RNA abundance and RNA synthesis (TED-seq/PRO-seq) is proportional to the RNA stability^15^. Using this ratio as the Stability Index (SI) is an approach to indirectly measure RNA half-life from RNA abundance and nascent transcription analysis. In HEK293 cells, SI was linearly correlated with direct RNA half-life measurement by 4-thiouridine (4sU) pulse chase^16^ (**Fig 2A**). We examined the various measures of SI using conventional RNA-seq, and 4sU half-life was best correlated when the SI was generated between TED-seq and PRO-seq (**Fig 2B, Fig S4**). Therefore, STOAT provides a prediction of RNA half-life without metabolic labeling or perturbation time-course.

**Fig 2.**
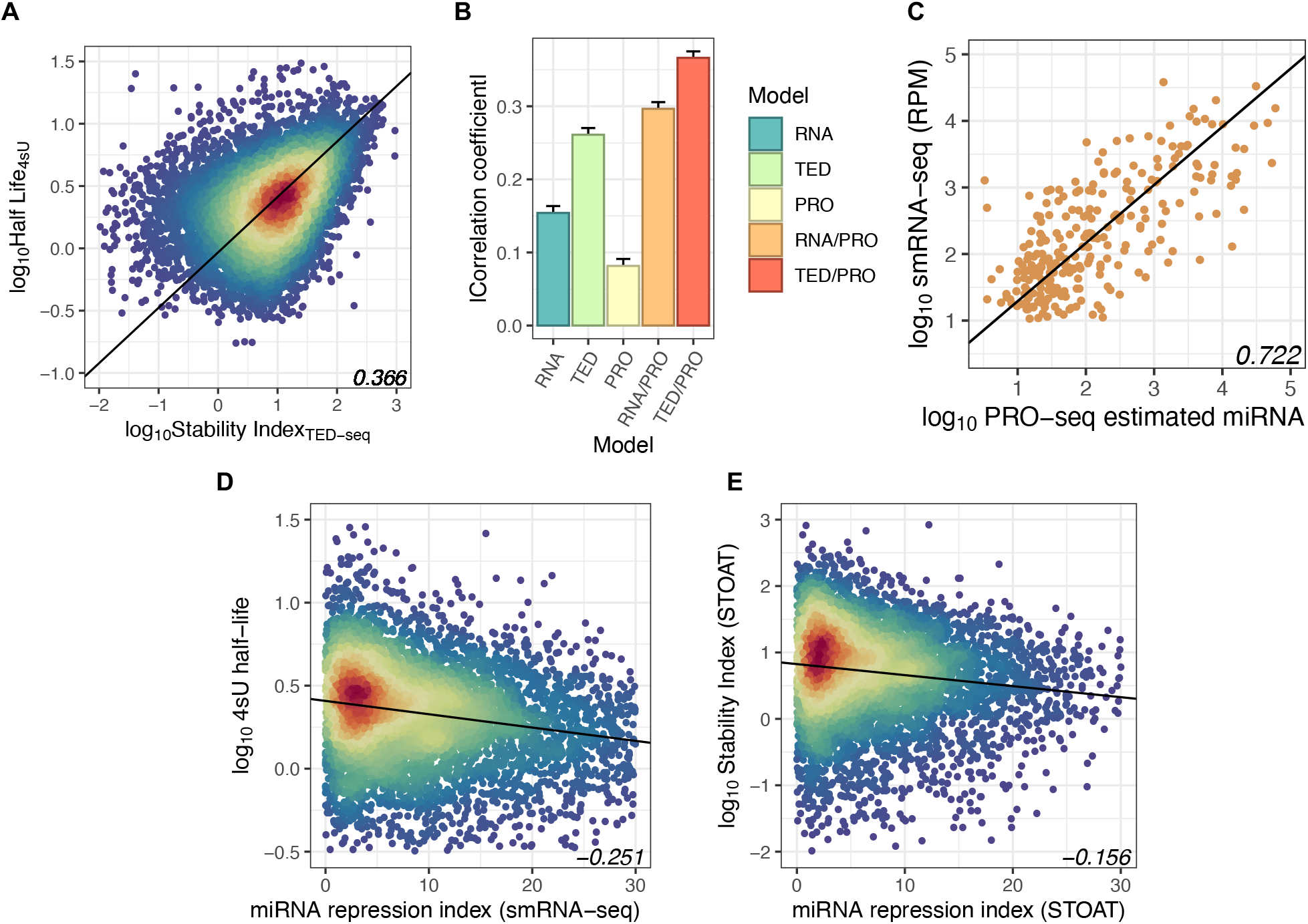
Reconstitution of RNA Stability and miRNA Network Using STOAT. **A**. Correlation between Stability Index using TED-seq and RNA stability measurements using 4sU labeling *(r= 0*.*366)*. **B**. Strengths of association between RNA stability (4sU) and expression measures and stability indices for RNA-seq, TED-seq, PRO-seq. **C**. Correlation between estimated miRNA expression from PRO-seq and smRNA-seq *(r= 0*.*722)*. **D**. Correlation between targets can be based on miRNA repression index and target mRNA stability using smRNA-seq and 4sU-seq. **E**. Correlation between miRNA repression index and mRNA stability index by STOAT (right).

Furthermore, STOAT was employed to dissect a post-transcriptional network. MicroRNAs (miRNAs) are a major post-transcriptional regulation mechanism^17,18^. While small RNA-seq is the standard assay, nascent transcription of miRNA genes also provides a reasonable estimate of miRNA expression^19^ (**Fig 2C, Supplementary Text, Fig S5, Table S2, S3**). We further investigated connecting the miRNA expression levels to their target mRNA regulation. We defined the miRNA Repression Index (miR-RI) of each mRNA based on TargetScan prediction and miRNA expression level. As expected, the miR-RI of target mRNAs based on smRNA-seq data in HEK293 cells was negatively correlated with target 4sU stability data (**Fig 2D**). We also observed similar negative correlation using PRO-seq-based miRNA estimates and SI (**Fig 2E**), indicating that miRNA post-transcriptional networks can be dissected using STOAT without small RNA or metabolic RNA labeling data.

### Cell Type Specific Analysis of Poly(A) Tail Lengths

Next, we investigated the global and cell type-specific differences in poly(A) tail lengths (PAL). While the classical view of polyadenylation is that it promotes mRNA stability and translation, recent studies show that longer PAL is associated with lower mRNA stability and translation, suggesting PAL reaches optimal size from poly(A) tail pruning ^20-23^. Still, the question remains: why does variability of PALs exist across transcripts and cell types? Transcriptome-wide poly(A) tail methods such as PAL-seq ^24^ or TAIL-seq ^25^ provide high quality PAL measurements, but they require modification to standard sequencing devices. Long read mRNA sequencing also provides information about PAL but is currently not cost efficient ^26-30^. To address this question, we used our previously developed TED-seq, which is cost-efficient and compatible with standard sequencing procedures for higher throughput PAL measurements ^14,15^.

By applying this tool, we found that global PALs were distributed between 50 to 100 bases (**Fig S6**). Non-coding RNAs (ncRNAs), such as long non-coding RNAs (lncRNA), and newly annotated ncRNAs (RP-ncRNA), which include enhancer RNAs (eRNAs) and upstream antisense RNAs (uaRNAs), overall showed longer PAL distributions than mRNAs (**Fig 3A**). This is compatible with previous studies reporting that actively translated RNAs have shorter PALs, demonstrating the advantage of TED-seq.

**Fig 3.**
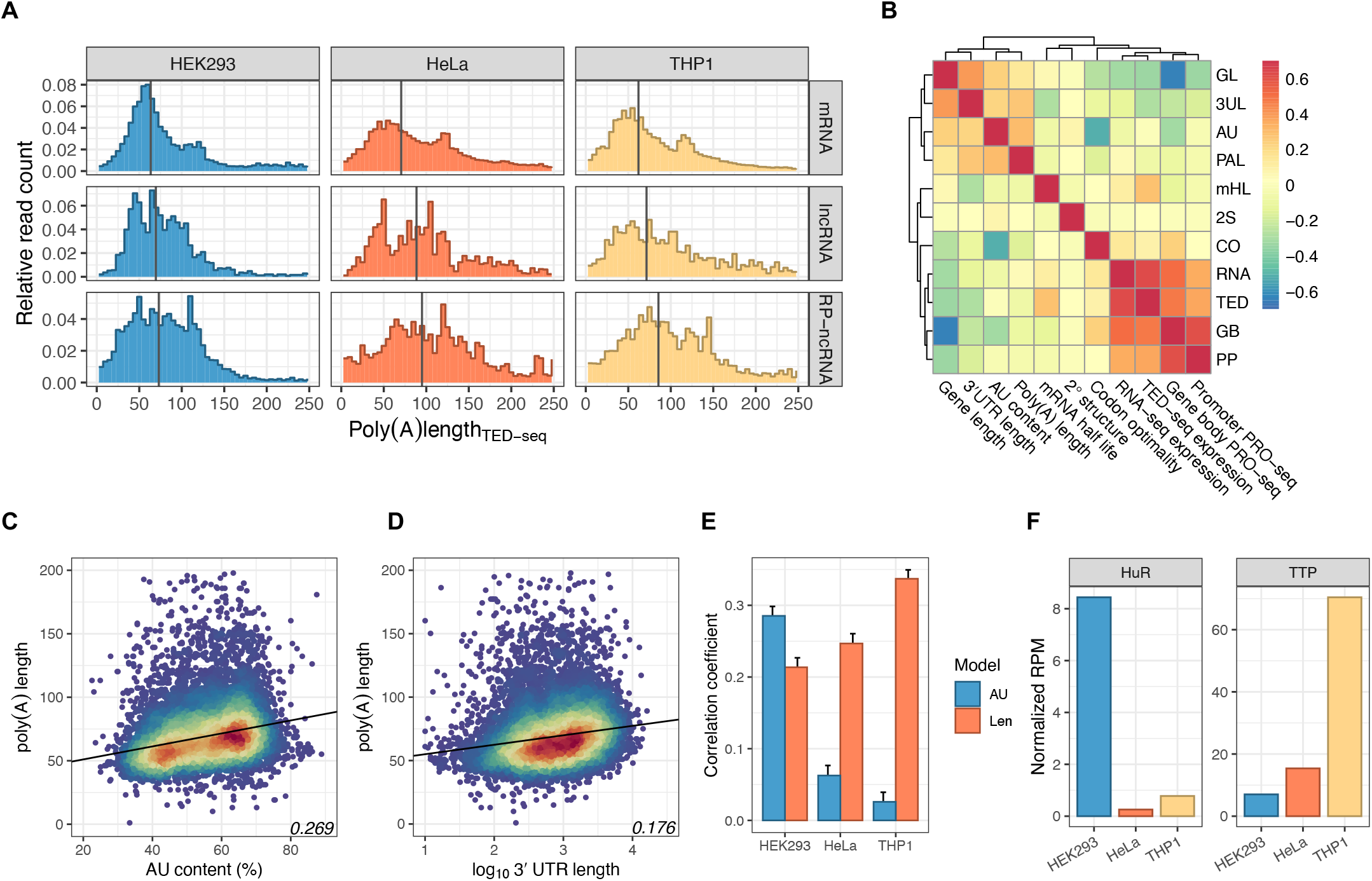
Cell Type Specific Analysis of Poly(A) Tail Lengths. **A**. Distribution of poly(A) lengths in all RNA molecules in different human cell lines, HEK293, HeLa and THP1 in mRNA, IncRNA, and RP-ncRNA. **B**. Heatmap of PAL correlation with mRNA and transcription features in HEK293 cell. TED-seq expression for mHL (mRNA half-life) and PAL length shows higher correlation than that of RNA-seq itself. **C**. Scatterplot of correlation between PAL and AU content with PAL. **D**. Scatterplot of correlation between PAL and 3′ UTR length with PAL. **E**. Correlation between PAL (Len) and AU content or 3′ UTR length across different cell types. **F**. RPM expression levels of HuR and TTP across different cell types quantified by TED-seq.

We further correlated PAL with various features of the transcripts and genes (**Fig 3B**). Among the features we tested, PAL was most strongly correlated with AU content (**Fig 3C**) and the length of the 3′ UTRs (**Fig 3D**). These closely correlated features were RNA sequence features rather than RNA expression features. While HEK293 cells had stronger correlation between PAL and AU content, other cell types we surveyed showed stronger correlation to 3′ UTR lengths (**Fig 3E, Fig S7**). This suggests that there is a cell type-specific PAL regulatory mechanism that depends on sequence elements and AU rich element (ARE) RNA binding proteins. Some of the examples of ARE binding proteins that stabilize or destabilize target transcripts are HuR and tristetraprolin (TTP), respectively. We found that, in HEK293 cells, PAL was more strongly associated with ARE, and HuR was more predominantly expressed (**Fig 3F**). This supports the hypothesis that the stabilizing effect of HuR mediates the association between AREs to longer PALs. In other cell types, the HuR-TTP balance also coincides with ARE-PAL association (**Fig 3F**), suggesting that HuR-TTP balance links PAL and transcript stability.

### Characterization of 3′ Processing in Novel Transcript Isoforms

Here we identified new transcript isoforms using the characteristic peak patterns of PRO-seq and TED-seq at 5′ and 3′ends of genes which allowed us to discover new transcript isoforms. Upon surveying the STOAT-based de novo transcription units, we noticed two types of new 3′ ends. One was short TSS-associated polyadenylated transcripts, which may arise from uaRNA or intronic transcription (**Supplementary text, Fig S8**). We observed that their PALs were comparable to mature transcripts, suggesting that they use the same cleavage-polyadenylation machinery as mRNAs.

The second finding was the prevalence of multiple alternative CPSs within long 3′ UTRs which were often un-annotated. For example, we identified three different alternative 3′ CPS sites for a connective tissue disease-related gene, B3GALT6^31^, two of which have not been previously annotated (**Fig 4A**). The RNA-seq base annotation only identified a single exon transcript, and the other 2 isoforms were detected through TED-seq. By comparing TED-seq with RNA-seq, the prevalence of alternative 3′ CPS was transcriptome wide. On average, ∼0.6 alternative 3′ CPSs were discovered per 3′ UTR using TED-seq, compared to ∼0.2 GENCODE annotated alternative 3′ CPS per RNA-seq 3′ UTR (**Fig 4B**).

**Fig 4.**
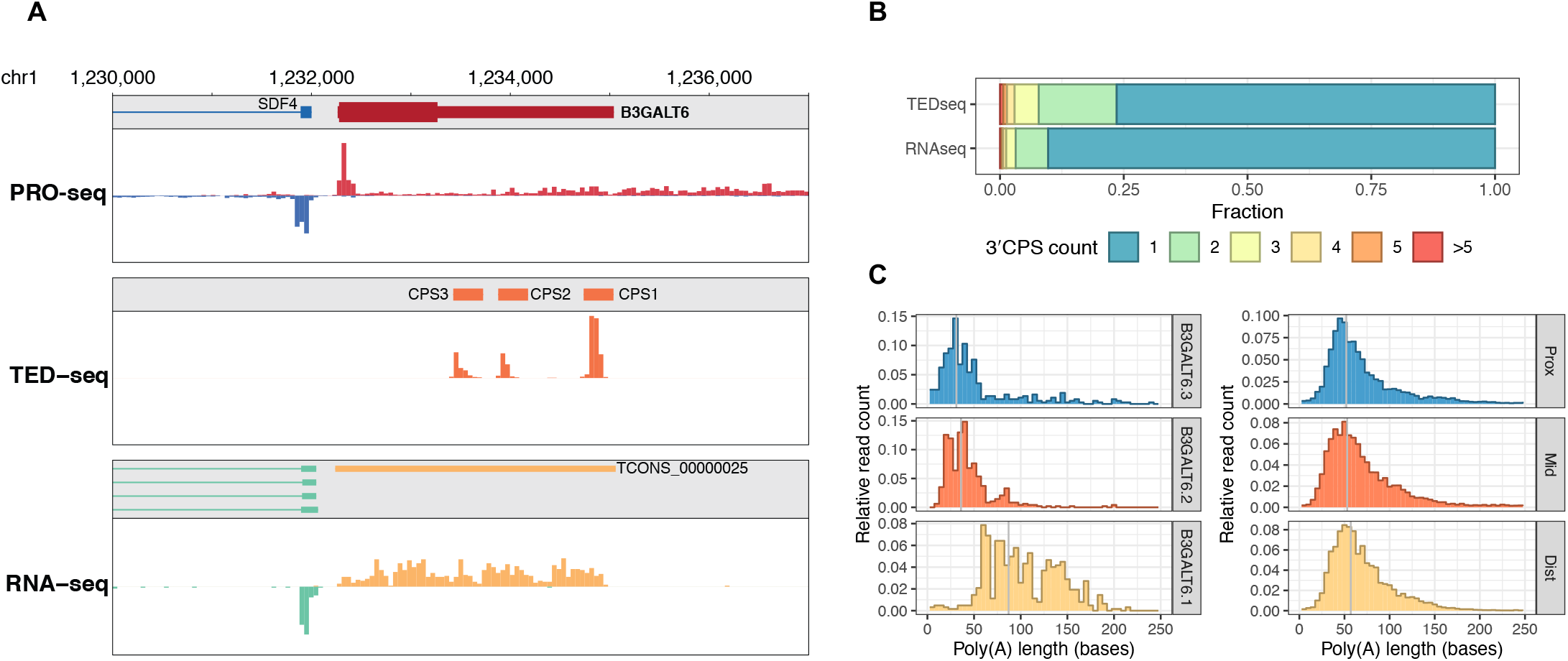
Characterization of 3′ Processing in Novel Transcript Isoforms. **A**. Discovering unannotated novel polyadenylated transcripts, 3′ CPS isoforms, in B3GALT6 gene. **B**. Finding number of alternative polyadenylation sites, 3′ CPS, discovered by STOAT **C**. Distribution of PAL differences in B3GALT6 isoforms (left), and average PAL profiling in alternative 3′ CPS isoforms (right).

These multiple alternative 3′ CPS sites showed isoform-dependent differential poly(A) lengths. For example, poly(A) tail length distributions of the three B3GALT6 3′ end isoforms all differed, the longest isoform had the longest poly(A) tails (**Fig 4C, left**). Globally, alternative 3′ ends that were the most distal showed modest but consistently longer PALs (**Fig 4C, right**). This agrees with the global observation that poly(A) tail lengths correlate with 3′UTR lengths and suggests that a 3′ UTR-dependent regulation of polyadenylation may take place that discriminates against certain isoforms^24-25, 32-34^. It is also possible that longer poly(A) tails could be a compensatory mechanism to counteract reduced stability. Alternatively, shorter poly(A) tails may be the consequence of more stable short UTR transcripts undergoing gradual deadenylation.

## DISCUSSION

In summary, STOAT provides a streamlined solution that is cost-effective and scalable to generate a large number of measurements of RNA dynamics and processing. It provides an efficient way to estimate RNA stability through the combined analysis of RNA expression and PRO-seq. Estimates of miRNA expression can be linked to the mRNA stability to elucidate post-transcriptional regulatory networks in different cell types with minimal cost. Cell type-specific transcription and polyadenylation reveal mRNA features that differentially affect RNA dynamics and processing. By integrating the analysis of RNA synthesis and processing, STOAT provides an added dimension of RNA dynamics, stability, and post-transcriptional regulatory networks.

Conventional measures of mRNA stability analysis required dense time-course measurements either after transcriptional inhibition or metabolic RNA labeling, which is laborious and may not be applicable in tissue samples. STOAT serves as an effective alternative measure of these quantities of RNA dynamics. Poly(A) tail length measurements are additional quantities related to post-transcriptional RNA processing. Compared to other poly(A) tail methods, TED-seq has proven to be cost-efficient and compatible with standard sequencing procedures that can be more widely used. Overall, STOAT provides a realistic, alternative, streamlined solution to generate large scale measurements of RNA dynamics and processing.

Besides being an efficient tool for RNA expression and stability, STOAT also provided novel insights by further dissecting miRNA networks and poly(A) length (PAL) regulation. miRNA is one of the most important factors of post-transcriptional regulation. While small RNA-seq is the standard assay, measuring transcription at miRNA loci by PRO-seq also provided estimates of miRNA expression levels. We correlated miRNA expression to the repression of target mRNA repression to constitute miRNA regulatory networks within the minimal STOAT framework that are specific to different human cell lines (**Supplementary Text**).

PAL is another major RNA modification associated with RNA stability and translation under rapid transient post-transcriptional responses, such as early development, cell cycle, and stress response ^24,32-34^. Our correlative analysis of PAL with various features of the transcriptome across cell types suggests that information directing PAL is encoded in the sequence, such as the AU content and 3′ UTR length; the encoded information is variably interpreted across cell types. Cell type-specific dependency of PAL on AU rich elements (AREs) may be a consequence of variable expression of ARE binding proteins such as HuR and TTP. This variable expression has a spectrum of effects on mRNA, ranging between stabilization and destabilization.

Overall, STOAT reveals various aspects of gene transcriptional regulations simultaneously. We propose that this combination of PRO-seq and TED-seq has potential to be used as a minimal set of gene expression analysis that provides a stereoscopic view of the dynamics of transcriptome.

## LIMITATIONS OF STUDY

For STOAT to perform successfully, at least a pair of high-throughput dataset is necessary; a nascent RNA-sequencing and a RNA-seq from the same cell-type and condition. Preferable datasets for STOAT are PRO-seq^12,13^ and TED-seq^14,15^. However, other types of poly(A) tail length sequencing data such as TAIL-seq or PAL-seq can replace TED-seq to utilize the full functionality of STOAT. Instead of TED-seq, other 3’-RNA-seq based methods can be used for the demonstrated analyses except for the poly(A) tail analysis. Other forms of nascent RNA-seq data such as GRO-seq, 4sU-seq can be used instead of PRO-seq but with decreased resolution and precision. Whereas current version of STOAT is optimized for PRO-seq and TED-seq, it can be used for various applications for dynamics analyses when de novo annotation, RNA stability, and miRNA analyses are required while only steady-state datasets are available.

## ACKNOWLEDGEMENTS

We thank the current and previous members of the Kwak lab and the Department of Molecular Biology and Genetics at Cornell University for providing constructive discussion and sharing unpublished datasets for this study. This study was supported by NIH 1R35GM142979 and discretionary funds (Cornell University) to HK.

## AUTHOR CONTRIBUTIONS

SAL & HK performed computational analysis and wrote the manuscript. All authors read and approved the final manuscript.

## INCLUSION AND DIVERSITY STATEMENT

Our dataset includes the use of HeLa cells, a cell line from an African American patient Henrietta Lacks. While the inclusion of this cell line was not initially intended for diversifying our dataset, we acknowledge that the use of HeLa cells has greatly contributed to biomedical science including our own study. We also acknowledge that the acquisition and shared use of HeLa cells was a result of historically exploited minorities and will promote to diversify genomic data inclusion for medically underrepresented minorities. We support inclusive, diverse, and equitable conduct of research.

## STAR METHODS

### Resource availability

#### Lead contact

Further information and requests for resources should be directed to the lead contact, Hojoong Kwak.

#### Materials availability

This study did not generate new unique reagents.

#### Data and Code Availability

We used previously published RNA-seq, PRO-seq, and Tail End Displacement sequencing (TED-seq) data in HEK293, HeLa, and THP1 cells by Woo et al and Kwak et al.^14^, and re-analyzed the raw sequencing data (GEO: GSE103719, GSE161188). Scripts for the stereoscopic analysis of the transcriptome (stoat) are available in github (https://github.com/sl2665/stoat).

This paper does not report original code. Any additional information required to reanalyze the data reported in this paper is available from the lead contact upon request.

### Method details

#### Quantification of PRO-seq, TED-seq, and RNA-seq expression

Quantification of the PRO-seq data was performed as described previously ^13^. Briefly, we used 250 bp upstream and down-stream of the TSS (500 bp region) as the promoter region to calculate promoter read counts. For gene body counts, we used the base coverage collapsing multiple reads at single positions to one read to avoid overcounting gene body reads. Both the promoter and gene body PRO-seq counts were normalized to reads/fragments per kilobase per million mapped reads (RPKM/FPKM). For the RNA-seq analysis, we used cufflinks ^36^ with Gencode V26 annotation as a reference to calculate isoform specific FPKM levels in our RNA-seq data. For the quantification of TED-seq data, we generated the last 500 bases regions of the Gencode V26 annotations from the 3′ CPS and quantified TED-seq read count in FPKM in the last 500 bases regions.

#### TED-seq poly(A) tail length processing pipeline

We used TED-seq to precisely measure the poly(A) tail lengths (PAL) of transcripts aligned to the whole genome which is compatible with splice junctions and novel transcriptome annotations. For precise mapping of the 5′ ends of TED-seq reads spanning through potential splice junction, the UMI collapsed aligned reads (bam format) were separated and pooled by the number of soft-clipped bases at the 5′ end of the reads.

The soft-clipped bases are derived from short portions of the sequence reads spanning over a splice junction but not long enough to be mapped to the upstream exon. We then calculated the distribution of the 5′ ends of TED-seq reads up to 500 bases from the 3′ CPS in the exons, concatenating any upstream exons within 500 bases of the 3′ CPS. The TED-seq reads with soft-clipped 5′ ends were shifted upstream by the number of soft-clipped bases and added up to the distributions. Finally, the TED-seq reads distributions are shifted downstream by the insert size of the library (300 base) to compensate for the displacement made by library size selection (**Fig S1A**). A total of 4 TED-seq replicates were used to produce PAL distributions for annotated transcripts.

#### Differential expression and poly(A) tail length analysis

We used raw read counts and DESeq2 to identify differentially expressed genes in PRO-seq and TED-seq. For RNA-seq, we used Cufflinks generated adjusted p-values to find differentially expressed genes. To identify transcripts with differential poly(A) tail lengths, we used the pipeline above, and identified transcripts with the differences of median poly(A) tail lengths greater than 20 bases.

#### Calculation of Stability Index

We defined the Stability Index (SI) as TED-seq or RNA-seq RPKM divided by PRO-seq RPKM in each transcript. Transcripts with PRO-seq gene body RPKM greater than 0.1 and TED-seq/RNA-seq RPKM/FPKM greater than 1.0 were used.

#### miRNA network analysis

To measure miRNA transcription level using PRO-seq, we used Gen-code V26 annotations to define the transcription unit (annotated coding or non-coding RNA) that transcribes through the miRNA site in the same strand. We used the PRO-seq RPKM at the miRNA region 1 kb upstream and downstream to define the transcription level of the pri-miRNA transcript. The ratio between small RNA-seq (reads per million: RPM) / PRO-seq (RPKM at miRNA region) is defined as miRNA processing-stability index (PSI) for each miRNA. miRNA PSIs were calculated in both HEK293 and HeLa data, and average of the two were used as the universal miRNA PSI. The estimated miRNA expression level (RPM) in other cell types (e.g. THP1) was calculated by multiplying the universal PSI to PRO-seq RPKM at the 2 kb window at the miRNA region. miRNA repression index (miR-RI) was calculated as the sum of the log10 of the product of TargetScan binding scores and miRNA expression level in reads per million (RPM). Either small RNA-seq miRNA quantification or estimated RPM from PRO-seq data were used (See supplementary text, Quantification of miRNA Expression and Further Dissection of Its Regulatory Network).

#### Quantification of transcript features associated with PAL

Transcript features were based on GENCODE v26 annotation and hg38 reference genome sequence. Gene length and 3′ UTR lengths were extracted from the GENCODE gene annotation. Secondary structure scores are calculated from the 3′ UTR sequence and ViennaRNA RNAfold prediction scores. The 3′ UTR were divided into bins of 100 bases, and the average of the local m-fold folding energies of the 100 bases were calculated and used as the secondary structure scores. Codon optimality was calculated as described previously using the GENCODE coding sequence^35^.

#### Transcriptome assembly from the RNA-seq dataset

We removed the first 8mer unique molecular indices (UMIs) of the RNA-seq reads and tagged it to the read IDs followed by the alignment to the hg38 reference genome using STAR aligner. We then removed PCR duplicates by collapsing the reads with the same UMI mapped to the same genomic positions (total of 135.2 million uniquely mapped reads). We assembled the transcriptome using cufflinks with the strand-specific second stranded option and followed the standard procedure otherwise as described by Trapnell et al,^37^. Total of 4 replicates of strand specific RNA-seq dataset was used, and the transcriptome assemblies were merged using cuffmerge, yielding 24,950 genes with 69,498 transcript isoforms.

#### De novo annotation of 3′ CPS and alternative 3′ CPSs

To identify 3′ CPS de novo, we collected TED-seq reads that contain at least 15 consecutive A bases at the end of the reads, that may contain the junction between 3′ UTRs and poly(A) tails. These sequences were merged from all TED-seq raw data, and the poly(A) tail sequences were clipped using cutadapt. The clipped sequences were aligned to the hg38 reference genome using the STAR aligner. Genomic coordinates containing more than or equal to 10 reads, at which the 3′ ends of the aligned TED-seq sequences were positioned, were defined as de novo 3′ CPS. Then we defined alternative 3′ CPS isoforms as the TED-seq 3′ CPS that are within the last exons of Gencode V26 genes annotation and do not overlap with any Gencode 3′ end annotations within 100 bp. Short TSS associated polyadenylated transcripts were defined by first identifying Gencode 5′ TSSs (n = 178,292) that are expressed in HEK293 cells by having an active dREG site occupying the TSS (n = 66,026), then identifying TED-seq 3′ CPS within 5 kb or 2 kb from the TSSs on both strands (n = 2,715 or 1,109 respectively). We classified these short transcription units based on their orientation from the TSS and overlap with the Gencode annotations into novel intragenic or extragenic transcription units. The extragenic transcription units on the antisense strand from its associated TSS were defined as polyadenylated uaRNAs. The intronic 3′ CPS were selected from the intragenic transcription units, and the full length 3′ CPS from the Gencode annotation defined as their host genes for the intronic 3′ CPSs.

