## Supplementary figures and images for "Stereoscopic RNA Analysis of Transcriptional and Post-transcriptional Regulation"

### Sup

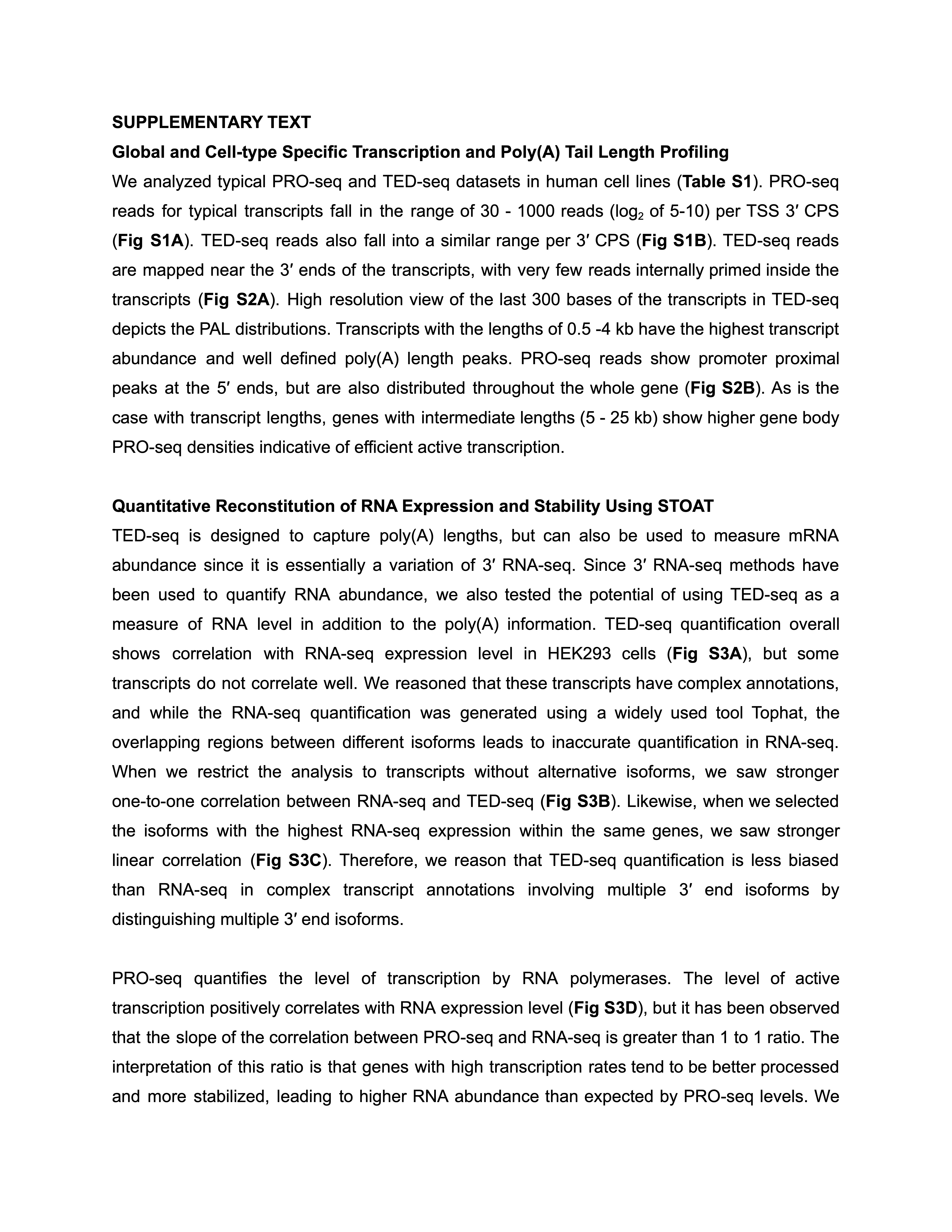
